# Transcriptomic landscape of sex differences in obesity and type 2 diabetes in subcutaneous adipose tissue

**DOI:** 10.1101/2024.09.18.613678

**Authors:** Roxana Andreea Moldovan, Marta R. Hidalgo, Helena Castañé, Andrea Jiménez-Franco, Jorge Joven, Deborah J. Burks, Amparo Galán, Francisco García-García

## Abstract

Obesity represents a significant risk factor in the development of type 2 diabetes (T2D), a chronic metabolic disorder characterized by elevated blood glucose levels. Significant sex differences have been identified in the prevalence, development, and pathophysiology of obesity and T2D; however, the underlying molecular mechanisms remain unclear. This study aims to identify sex-specific biomarkers in obesity and T2D and enhance our understanding of the underlying mechanisms associated with sex differences by integrating expression data.

A systematic review, individual transcriptomic analysis, gene-level meta-analysis, and functional characterization were performed to achieve this aim. Eight studies and 236 subcutaneous adipose tissue samples were analyzed, identifying common and sex-specific biomarkers, many of which were previously associated with obesity or T2D.

The obesity meta-analysis yielded nineteen differentially-expressed biomarkers from a sex-specific perspective (e.g., *SPATA18, KREMEN1, NPY4R*, and *PRM3*), while a comparison of the expression profiles between sexes in T2D prompted the identification and validation of specific transcriptomic signatures in males (*SAMD9, NBPF3, LDHD*, and *EHD3*) and females (*RETN, HEY1, PLPP2*, and *PM20D2*). At the functional level, we highlighted the fundamental role of the Wnt pathway in the development of obesity and T2D in females and the roles of more significant mitochondrial damage and free fatty acids in males.

Overall, our sex-specific meta-analyses supported the detection of differentially expressed genes in males and females associated with the development of obesity and T2D, emphasizing the relevance of sex-based information in biomedical data and opening new avenues for research.

**Highlights:**

- First meta-analysis on WAT with sex as a central perspective in obesity and T2D
- This study identifies 19 sex-differential biomarkers in obesity, highlighting NPY4
- Sex specific transcriptional signatures in SAT in the development of T2D
- Wnt pathway genes show sex-specific roles in obesity and T2D, notably in females
- Obesity increases mitogenesis in male, mediated by SPATA18, with an increased role of free fatty acids in T2D

## Introduction

Diabetes mellitus is a chronic metabolic disorder characterized by elevated blood glucose levels due to insulin resistance or insulin absence [1]. Depending on disease etiology, diabetes has been classified into several categories, with type 2 diabetes (T2D) being the most relevant [2]. T2D is caused by the dysregulation of carbohydrate, lipid, and protein metabolism due to insulin resistance at the peripheral level and simultaneous pancreatic β-cell dysfunction [3] and accounts for over 90% of global diabetes cases. While T2D has a minor genetic component, lifestyle, age, and weight have substantial influencing roles [4].

T2D’s impact represents a global health concern; in 2021, the International Diabetes Federation estimated that ∼537 million individuals aged 20-79 were affected, representing ∼9.8% of the world’s population. This percentage increased in older age groups, surpassing 20% in individuals over 70 [5]. Epidemiological studies attribute this escalating trend to lifestyle changes, with obesity emerging as a primary contributor [6, 7].

Obesity - a chronic, multifactorial, and treatable disorder of neurobehavioral origin - is characterized by increased body fat, leading to dysfunction in adipose tissue, which plays a crucial role in metabolic homeostasis and severe adverse effects on metabolic, biomechanical, and psychosocial health [8, 9]. The literature describes two main types of adipose tissue based on their structural characteristics, coloring, vascularization, and function: white adipose tissue (WAT) and brown adipose tissue [10, 11].

WAT stores energy in the form of triglycerides, and its quantity increases with age and body weight. Under normal conditions, insulin acts on WAT via the PI3K/AKT signaling pathway, stimulating adipogenesis, inhibiting lipolysis, and increasing fatty acid and glucose uptake [12, 13]. In pathological conditions such as obesity, adipose tissue dysregulation prompts the release of fatty acids and their metabolic intermediates into the bloodstream, which results in increased β-oxidation [14], oxidative stress [15, 16], and the secretion of pro-inflammatory hormones and cytokines [17, 18]. These factors significantly contribute to the development of insulin resistance and T2D via multiple signaling pathways, including PI3K/AKT [19, 20], RAS/MAPK [21–23], or Wnt [24–27]. Furthermore, inflammation can initiate diabetes by inducing changes in insulin sensitivity [28].

WAT is divided into two main deposits: visceral adipose tissue (VAT) and subcutaneous adipose tissue (SAT) [11, 17]. The impact of body fat levels and their distribution on insulin sensitivity remains critical [29]. Marked differences in gene expression patterns related to T2D exist between VAT and SAT, with VAT exhibiting a more pronounced and characteristic immunological pattern even in early disease stages [30, 31]. Conversely, SAT suffers from a less pronounced immune and inflammatory response and overexpresses genes related to the RAS/MAPK pathway and downregulates the expression of Peroxisome proliferator-activated receptor gamma (PPAR-γ), leading to altered adipogenesis [32–34]. These factors substantially increase T2D risk, positioning SAT not only as an early T2D marker, but also as a T2D risk marker, and a crucial nexus between T2D and obesity despite being less well-characterized than VAT [35].

The late consideration of sex in studies and the search for unified criteria for the diagnosis and treatment of diseases has led to ignoring differences at the physiological level and its role as a critical modifier in the factors surrounding a large number of diseases, including diabetes [36].

In relation to our focus, fundamental aspects of metabolic homeostasis display differential regulation between males and females [37, 38]. Sexual dimorphism, an evolutionary paradigm where females exhibit more significant resistance to the loss of energy reserves in the form of fat, plays a crucial role in metabolic homeostasis, obesity, and T2D [39]. The most significant sexual differences relate to genetic sex - the prenatal testosterone programming effect in males and the activating role of sex hormones during puberty [37, 40–42] - contribute to sexual biases in body fat distribution [43, 44], body mass index, and obesity [45, 46], and in T2D prevalence [47], susceptibility [41], progression [48, 49], diagnosis [50], and treatment [51].

In this context, VAT constitutes the primary type of fat storage in males, while females accumulate SAT [52, 53]. Adipose tissue constitutes 12-15% of male body weight but 20-27% of female’s [54]; furthermore, diabetes prevalence in females aged 20-79 remains slightly lower than in males (10.2% vs. 10.8%) [5]. While these differences are clinically understood, the underlying molecular mechanisms remain unknown; thus, this study aimed to characterize sex differences associated with obesity and T2D at the genic and functional level by integrating gene expression data of SAT and VAT available in public repositories. The results obtained in this work suggest a male- and female-specific SAT transcriptional signature in the development of obesity and T2D.

## Materials and methods

### Systematic review and study selection

A comprehensive systematic review, following the PRISMA guidelines [55, 56], was conducted during March 2023 to identify transcriptomics studies available in the Gene Expression Omnibus (GEO) and ArrayExpress databases (period: 2002-2022) [57–59] and the PubMed bibliographic search engine [60]. This search was based on the following inclusion criteria: i) keywords: “obesity,” “diabetes 2,” “type 2 diabetes,” “insulin resistance”; ii) organism: “Homo sapiens” or “Mus musculus”; iii) study type: “expression profiling by array” or “expression profiling by high throughput sequencing”; iv) sample count: ≤ 12 samples; v) tissue: “adipose tissue”; and vi) sex information: “gender” or “sex.” Subsequently, studies were filtered based on predefined exclusion criteria: i) experimental designs different from the comparison of cases and controls of obese and non-obese patients and/or insulin-sensitive and insulin-resistant individuals; ii) absence of information or representation of both sexes; iii) insufficient sample size (less than two samples per sex and group); and iv) tissue other than adipose.

### Bioinformatic analysis

The *in-silico* analysis employed a methodology comprising five steps: i) data download and preprocessing; ii) differential gene expression analysis (DGE); iii) DGE results integration using a meta-analysis-based approach; and iv) functional analysis. Phases i-ii were executed independently for each study, while phases iii-iv were dedicated to integrating and obtaining global results. All bioinformatics and statistical analysis were conducted using version 4.2.0 of the R software [61]. **Table S1** lists the packages used and their version numbers.

### Data download and preprocessing

Data from the selected studies were retrieved using the GEOquery [62] and ArrayExpress [63] packages. Normalized data was acquired whenever available, and the original authors’ normalization methods for each dataset were assessed. For microarray datasets, background correction, quantile normalization, and log2 transformation were performed using the affy [64] package when necessary, while RNA-seq data was converted to log2-counts-per-million using limma-voom [65]. All probe sets were annotated to NCBI Gene ID [66] using the AnnotationDbi package [67]. In the event of duplicate probe-to-gene mappings, the median expression was calculated. Furthermore, HUGO gene symbols [68] were included to aid further interpretation.

The study’s phenotypic variables were harmonized by categorizing patients and samples by sex (M for male and F for female), phenotype (Control for normal weight; Obesity for healthy obese; and T2D for type 2 diabetes obese), and tissue (SAT for subcutaneous adipose tissue; and VAT for visceral adipose tissue).

Exploratory analyses were conducted to detect patterns among the samples and the phenotypes of interest (group, sex, or tissue) and identify potential anomalies or batch effects. The analysis included box plots, hierarchical clustering based on Pearson correlation and Euclidean distance, and principal component analysis (PCA). Any observed batch effects in the samples were corrected using the sva package [69].

### Differential gene expression analysis

Differential gene expression analysis was conducted to study the development of obesity and T2D via two independent types of comparisons. In terms of obesity, the impact of obesity (IO) was examined by comparing obese patients and controls (Ob - C); the impact on males (IOM) was examined by comparing male obese patients and male control patients (Ob.M - C.M), and the impact on females (IOF) was examined by comparing female obese patients and female control patients (Ob.F - C.F). We also evaluated the sex-differential impact of obesity (SDIO) by comparing the two previous comparisons (Ob.M - C.M) - (Ob.F - C.F).

In the case of T2D development, the impact of T2D (ID) was examined by comparing obese patients with T2D and obese non-diabetic patients (T2D - Ob). A comparison between T2D obese males and obese insulin-sensitive males (T2D.M - Ob.M) was used to examine the impact of T2D on males, while a comparison between obese females with T2D and obese insulin-sensitive females (T2D.F - Ob.F) was used to examine the impact of T2D on females. The sex-differential impact of T2D (SDID) was also analyzed, representing the overall comparison of the two previous contrasts.

DGE was performed for each of the eight comparisons using the limma package [65]. The differential expression statistics were then calculated, and the p-values were adjusted for multiple comparisons using the Benjamini & Hochberg (BH) method [70]. A significance cut-off was established at a false discovery rate (FDR) < 0.05.

### Meta-analysis and results integration

Integrating individual DGE results employed a meta-analysis for each comparison and gene using a random effects model. Only genes present in at least two studies were considered, and logFC was employed as the effect size, with the DerSimonian-Laird estimator used to measure heterogeneity. This comprehensive analysis was performed using the metafor package [67]. The results yielded a robust measure of expression across all studies and provided a BH-adjusted p-value [14]. Significant genes were identified by applying a threshold of FDR < 0.05. These genes were then assessed based on their previously established links to obesity and T2D using the OpenTargets database (23.02 release) [71].

### Functional characterization

To identify and associate the gene consensus profiles obtained in meta-analysis with specific biological processes (BP), a gene set enrichment analysis (GSEA) was performed. For each analysis, a ranked gene list based on the logFC was analyzed using the clusterProfiler package with the Gene Ontology (GO) BP database [73] as the reference. The p-values were adjusted using the BH method [70], and logarithm 2 of the odds ratio (logOR) was used to assess the statistical effect. The significance cut-off was set at FDR < 0.05. The resulting terms were grouped based on their similarity using the rrvgo package [74] due to the redundancy and hierarchical structure of the GO BP database. The purpose of this grouping was to streamline the interpretation and comparison of results across different comparisons.

### Experimental validation

#### Study Selection and Sample Processing

The study included a total of n =32 adult patients with severe obesity (BMI>35 kg/m2) who underwent bariatric surgery at Hospital Sant Joan (Reus, Spain). SAT biopsies were obtained from male obese (n=8), female obese (n=8), male obese T2D (n=8), and female obese T2D (n=8) patients. The study received ethical approval from the Ethics Committee of Hospital Universitari Sant Joan (Reus, Spain) (EPIMET083/2018), and all participants signed the informed consent.

Total RNA was extracted from starting 100 mg of abdominal fat using a combined protocol including Trizol (Invitrogen) and RNA NucleoSpin® (Macherey-Nagel GmbH & Co) Kit. First-strand synthesis was performed using PrimeScript™ RT Reagent Kit (Perfect Real Time) (Takara).

#### Gene Expression Analysis

Quantitative gene expression analysis was performed on 50 ng cDNA templates. Real time-PCR was conducted in a LightCycler 480 Instrument IIR (Roche) using TB Green® PreMix ExTaq™ (Tli RNaseH Plus) (Takara). **Table S2** details the genes selected, as well as the primer list. Relative quantification was performed using the ΔCt method. Statistical analyses were performed with GraphPad Prism 10 (GraphPad Software V 10.0). Results are expressed as the arithmetic mean ± the standard error of the mean, and Student’s t-test was used. Any observed differences were considered significant when: p-value

<0.05 (*), p-value <0.01 (**) and p-value <0.001 (***).

## Results

### Systematic review and study selection

During the initial stage, the search and review of the GEO and ArrayExpress databases identified 404 studies (82% unique records); however, the application of the exclusion criteria provided a final number of eight records. The search for PubMed bibliographic sources involved a review of 230 articles; only three met the defined criteria, coinciding with the studies extracted from the databases. **Fig. 1** details the number of records in each phase and the reasons for exclusion.

**Fig. 1.**
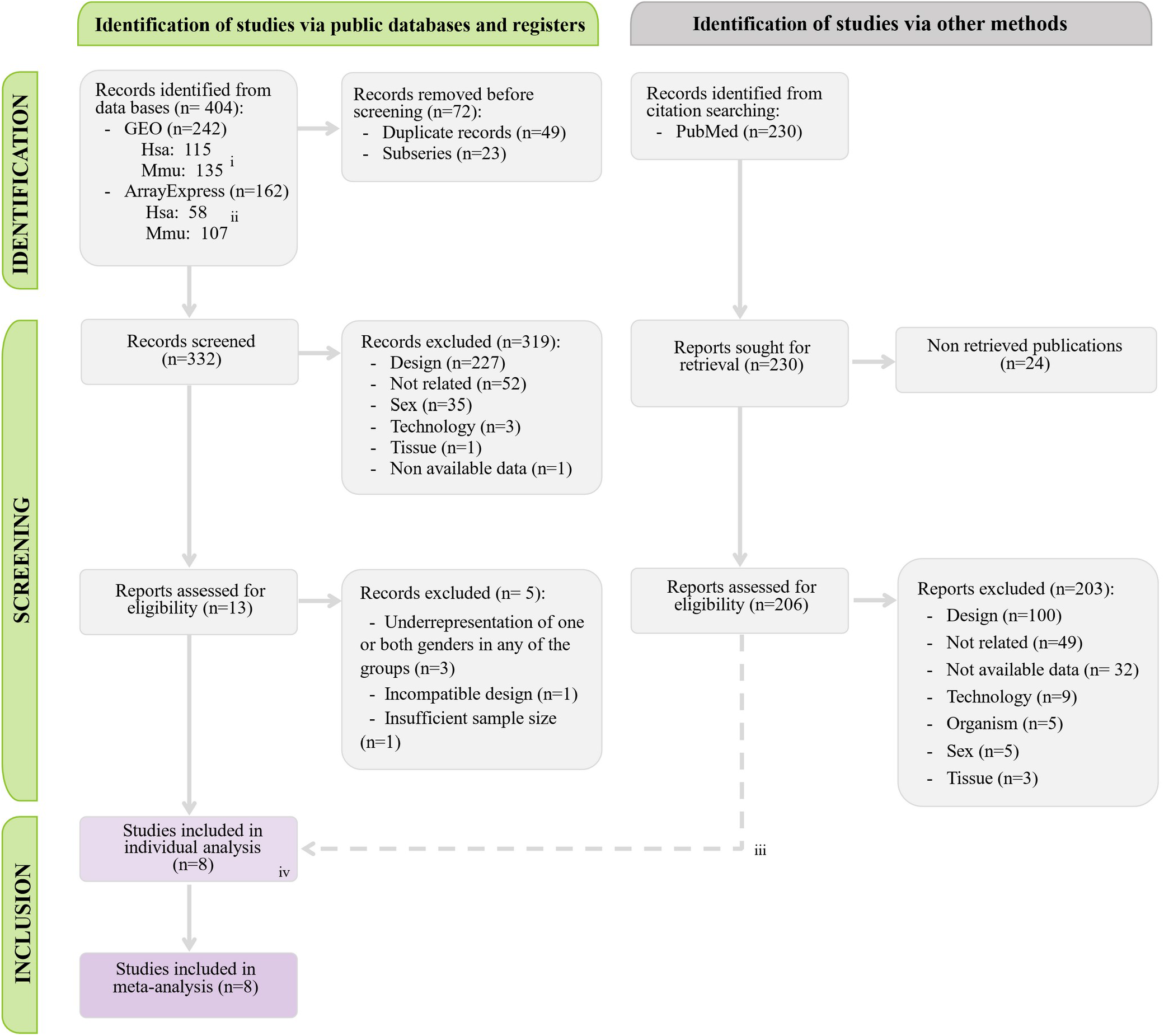
Flow diagram for the systematic review following 2020 PRISMA Statement guidelines. i) 8/242 studies included samples from both species; ii) 3/162 studies included samples from both species; iii) The three studies identified through bibliographic searching matched three records identified through database searches; iii) All samples included in the final study belong to *Homo sapiens*.

We selected eight studies (all *Homo sapiens*), which encompassed a total of 327 samples from three experimental groups (C, Ob, and T2D) and two tissues (VAT and SAT) (**Table 1**). Limitations in the number of studies, including VAT and their small sample size, prevented the individual analysis of this tissue. Furthermore, the dissimilarities noted between SAT and VAT during the exploratory analysis and as previously characterized [17, 30, 32, 34] led us to conclude that joint analysis would introduce noise into the meta-analysis. Consequently, we excluded the twenty samples corresponding to VAT. We analyzed 302 samples - 46.36% Control group; 48.01% Obese individuals; and 5.63% T2D patients – observing an overall sex distribution of 44.04% males and 55.96% females (**Table 2**). **Table S3** provides detailed information on the annotation process and the number of analyzed genes per study.

**Table 1.** Description of selected studies. Identifier accession in the database, database source, kind of platform, described phenotypes (C: control; Ob: healthy obese; T2D: type 2 diabetic obese), tissue (SAT: subcutaneous adipose tissue; VAT: visceral adipose tissue) and technology used are detailed for each study. All features can be found in the GEO and ArrayExpress databases.

| Study accession | Database | Platform | Phenotypes | Tissue | Type |
| --- | --- | --- | --- | --- | --- |
| E_MEXP_1425 [75] | ArrayExpress | Affymetrix GeneChip Human Genome U133 Plus 2.0 [HG-U133_Plus_2] | C, Ob | SAT | Microarray |
| GSE2508 [76, 77] | GEO | [HG_U95Av2] Affymetrix Human Genome U95 Version 2 Array<br>[HG_U95A] Affymetrix Human Genome U95A Array<br>[HG_U95B] Affymetrix Human Genome U95B Array<br>[HG_U95C] Affymetrix Human Genome U95C Array<br>[HG_U95D] Affymetrix Human Genome U95D Array<br>[HG_U95E] Affymetrix Human Genome U95E Array | C, Ob | SAT | Microarray |
| GSE20950 [78–80] | GEO | [HG-U133_Plus_2] Affymetrix Human Genome U133 Plus 2.0 Array | Ob, T2D | SAT & VAT | Microarray |
| GSE29718 [81] | GEO | [HuGene-1_0-st] Affymetrix Human Gene 1.0 ST Array [transcript (gene) version] | C, Ob | SAT & VAT | Microarray |
| GSE64567 [82] | GEO | Illumina HumanHT-12 V4.0 expression beadchip | C, Ob | SAT | Microarray |
| GSE92405 [83] | GEO | [HG-U133_Plus_2] Affymetrix Human Genome U133 Plus 2.0 Array [CDF: Brainarray HGU133Plus2_Hs_ENTREZG_v18] | C, Ob | SAT | Microarray |
| GSE141432 [84] | GEO | Illumina NextSeq 500 (Homo sapiens) | C, Ob, T2D | SAT | RNA-seq |
| GSE205668 [85] | GEO | Illumina HiSeq 2500 (Homo sapiens) | C, Ob | SAT | RNA-seq |

**Table 2.** Distribution of samples in studies by tissue, sex, and experimental group. C: control; Ob: insulin sensitive obese; T2D: type 2 diabetic obese; SAT: subcutaneous adipose tissue.

| Study | SAT |  |  |  |  |  | Total |
| --- | --- | --- | --- | --- | --- | --- | --- |
|  | C |  | Ob |  | T2D |  |  |
|  | M | F | M | F | M | F |  |
| E_MEXP_1425 | 8 | 5 | 9 | 5 | 0 | 0 | 27 |
| GSE2508 | 10 | 10 | 9 | 10 | 0 | 0 | 39 |
| GSE20950 | 0 | 0 | 2 | 8 | 4 | 5 | 19 |
| GSE29718 | 4 | 4 | 5 | 2 | 0 | 0 | 15 |
| GSE64567 | 10 | 19 | 9 | 26 | 0 | 0 | 64 |
| **GSE92405** | 9 | 17 | 9 | 17 | 0 | 0 | 52 |
| **GSE141432** | 3 | 6 | 4 | 4 | 4 | 4 | 25 |
| **GSE205668** | 22 | 13 | 12 | 14 | 0 | 0 | 61 |
| **Total** | 66 | 74 | 59 | 86 | 8 | 9 | 302 |

During the exploratory analysis phase, we identified anomalous clusters and correlations in samples from GSE2508, GSE20950, and GSE29718 caused by the use of two different platforms, preserving and processing samples at two separate times, and sequencing the samples in two batches. We corrected this batch effect during the individual analysis phase.

### Sex differences in obesity and T2D transcriptomic signatures

We performed two sets of four comparisons: IO, IOM, IOF, and SDIO for obesity, and ID, IDM, IDF, and SDID for T2D. The logFC indicates the magnitude of the change, while its sign indicates the direction of this change. Positive values indicate a higher expression in the first element of the comparison, while negative values would indicate an increase in the second.

In the differential expression analysis for obesity, we found significant differences in SDIO in 3/7 analyzed studies. Of note, study GSE29718 did not show significant differences in any comparisons. None of the individual studies demonstrated significant SDID comparison results in that context. **Table S4** describes the differential gene expression study-independent results.

We performed a meta-analysis for 37,226 genes in obesity and 20,161 genes in T2D, which included genes present in at least two independent studies (**Table 3**). Studying the impact of obesity in males and females, we observed the commonality of only 3.49% of genes in the IOM and IOF comparisons, comprising 22/428 upregulated genes and 4/300 downregulated genes based on the logFC (**Fig. 2A**). We observed the commonality of 7.64% of genes in T2D in the IDM and IDF comparison, comprising 49/639 upregulated genes and 66/982 downregulated genes (**Fig. 3** and **Table 3**).

**Table 3.** Meta-analysis results. Number of significant genes by comparison: IO (impact of obesity), IOM (impact of obesity in males), IOF (impact of obesity in females), SDIO (sex-differential impact of obesity), ID (impact of diabetes), IDM (impact of T2D in males), IDF (impact of T2D in females), SDID (sex-differential impact of T2D). Significant genes are separated by the sign of their logarithm 2 of fold change (logFC). The last row reports the number of genes included in each meta-analysis. The last row reports the number of genes included in each meta-analysis. The significant genes were identified when the adjusted p-value (False Discovery Rate) < 0.05.

|  | Obesity |  |  |  | T2D |  |  |  |
| --- | --- | --- | --- | --- | --- | --- | --- | --- |
|  | IO | IOM | IOF | SDIO | ID | IDM | IDF | SDID |
| **logFC > 0** | 430 | 277 | 147 | 10 | 713 | 105 | 534 | 0 |
| **logFC < 0** | 350 | 137 | 163 | 9 | 1,268 | 217 | 765 | 0 |
| **Total** | 780 | 414 | 310 | 19 | 1,981 | 322 | 1,299 | 0 |
| **Analyzed genes** | 37,226 |  |  |  | 20,161 |  |  |  |

**Fig. 2.**
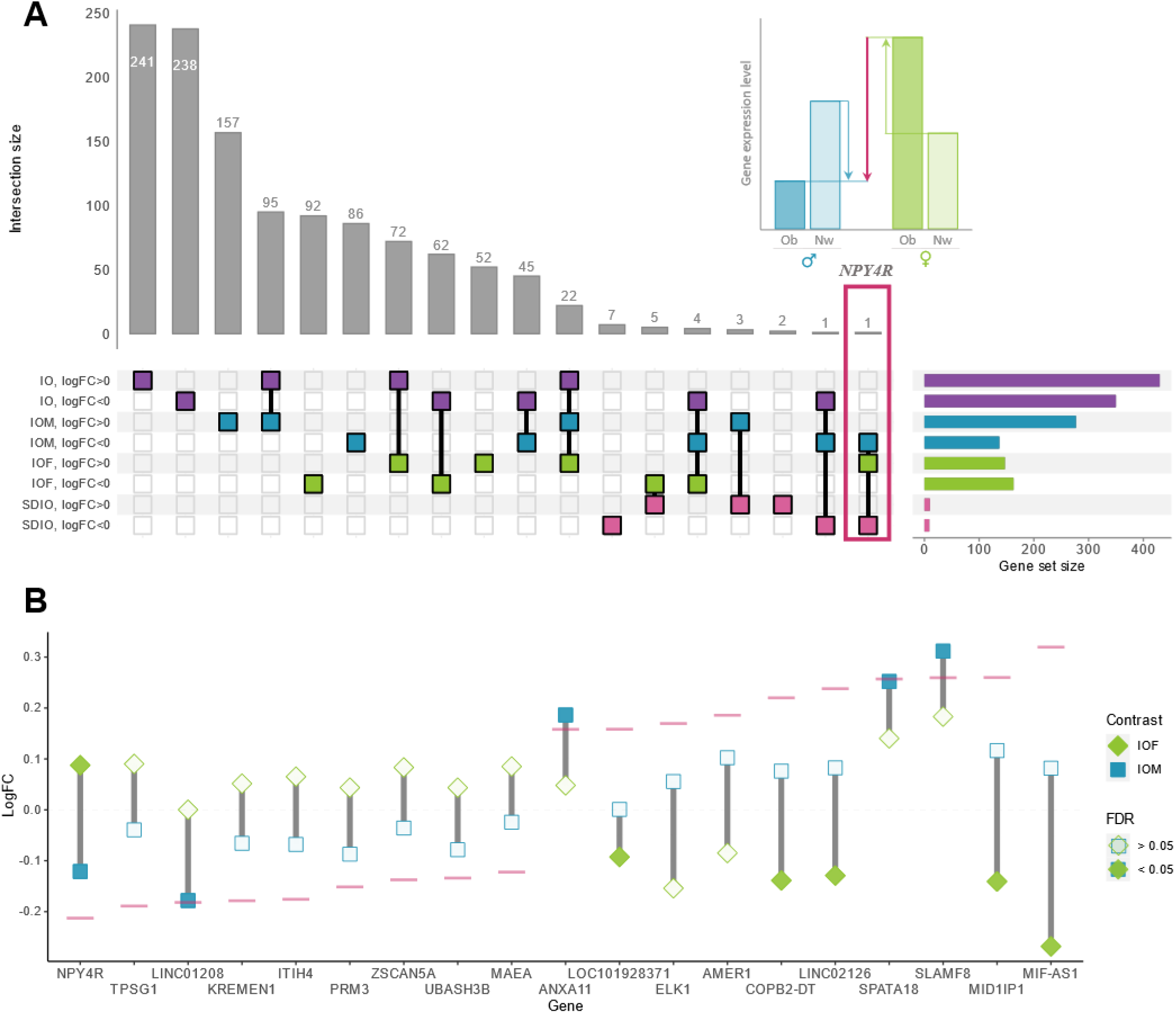
Transcriptomic profiles in the obesity meta-analysis. (**A**) Upset plot of significantly altered genes. Horizontally, the number of significant genes per comparison is displayed and separated by logFC. The purple, blue, green, and pink bars denote the IO, IOM, IOF, and SDIO comparisons, respectively, while the white and gray background represent positive and negative logFC values. The intersections between these groups are displayed vertically. The upper-right corner bar plot illustrates the variation of *NPY4R* expression levels. Blue and green arrows denote the variation in expression levels in the IOM and IOF comparisons, respectively, while the pink arrow indicates the overall effect of these changes in the SDIO comparison. (**B**) Exploration of the expression patterns of the nineteen differentially expressed genes in the SDIO comparison. Squares and rhombi represent the logFC obtained in the meta-analysis of IOM and IOF comparisons, respectively, while the pink line depicts the effect of these changes in the SDIO comparison. Significantly affected genes in the IOM and IOF comparisons are represented by filled squares and rhombi, respectively. Note that the mentioned colors refer to the online version of the article; please refer to the figure legend for further information. IOM: impact of obesity in males; IOF: impact of obesity in females; IDM: impact of T2D in males; IDF: impact of T2D in females.

**Fig. 3.**
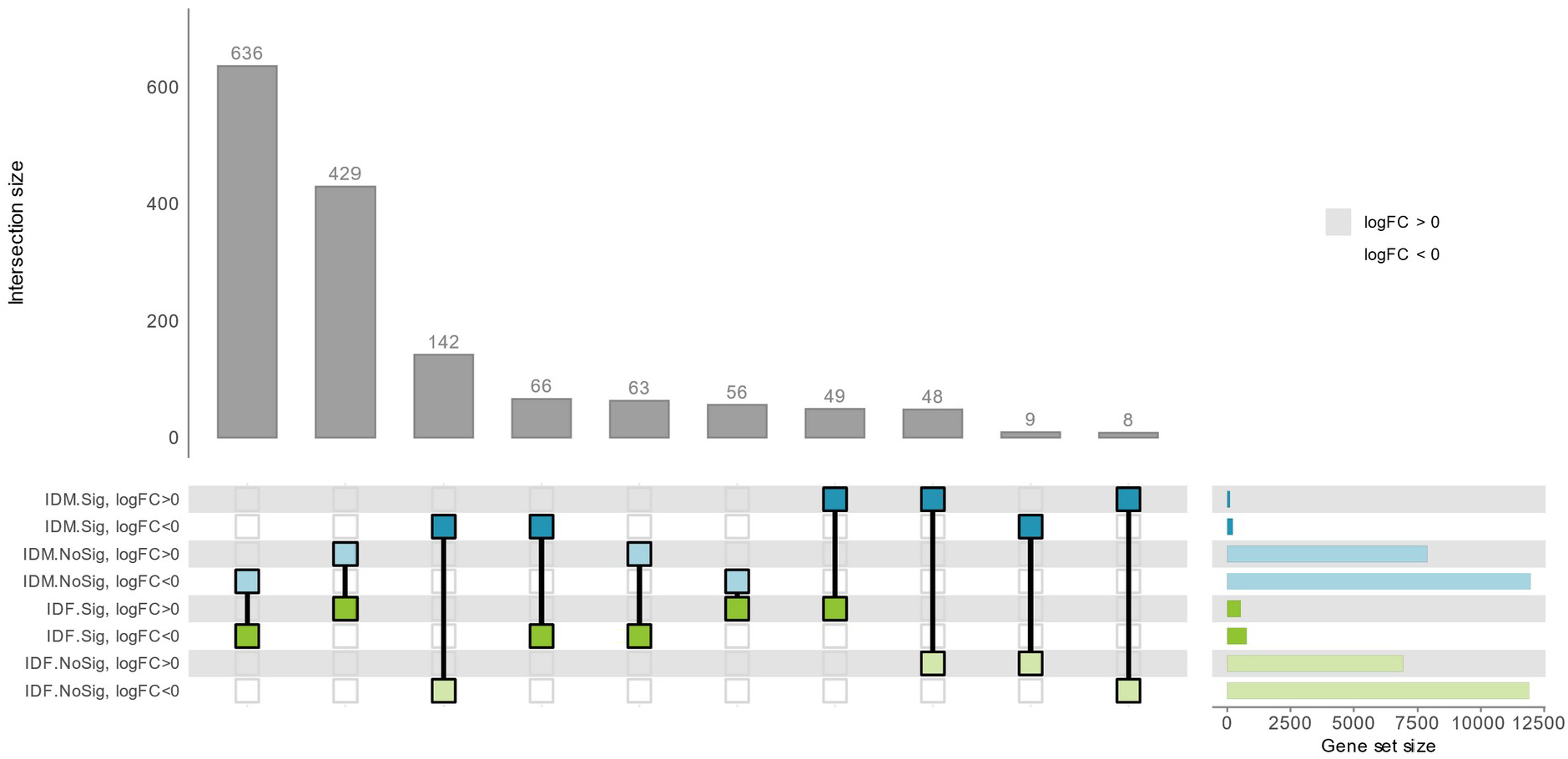
Transcriptomic profiles in the type 2 diabetes meta-analysis. An upset plot comparing expression profiles by sex was obtained in the IOM and IOF comparisons. Horizontally, the number of significant genes per comparison is displayed, separated by logFC and adjusted p-value. Intersections between these groups are displayed vertically. The IOM and IOF comparisons are denoted by blue and green color bars, respectively. Positive and negative logFC values are represented by white and gray backgrounds, with significance denoted by transparency. Significant values, using an adjusted p-value < 0.05, are opaque. Note that the mentioned colors refer to the online version of the article; please refer to the figure legend for further information. IOM: impact of obesity in males; IOF: impact of obesity in females; IDM: impact of T2D in males; IDF: impact of T2D in females.

### Transcriptomic profiles of sex differences in obesity

Exploring sex-specific differences in the impact of obesity (SDIO) led to the identification of nineteen differentially expressed genes as potential biomarkers. Comparative analysis of the SDIO results with previous comparisons provided valuable insight into the behavior of these genes during the development of obesity in males and females (**Fig. 2**).

The comparison reports differences in gene expression that may be associated with various physiological scenarios resulting from a combination of the IOM and IOF comparisons. **Fig. 2B** depicts the differences in the logFC of the differentially expressed genes in the SDIO comparison. 4/19 genes displayed an opposite pattern in the sex-specific comparisons. The *NPY4R, TPSG1, KREMEN1, ITIH4, PRM3, ZSCAN5A, UBASH3B*, and *MAEA* genes displayed upregulated expression in obese females and downregulated expression in obese males, whereas *ELK1, AMER1, COPB2-DT, LINC02126, MID1IP1*, and *MIF-AS1* displayed upregulated expression in obese males and downregulated expression in obese females. Of note, *LINC01208* and *LOC101928371* expression became significantly decreased exclusively in one sex, while *ANXA11, SPATA18*, and *SLAMF8* expression underwent a concordant increase in both comparisons with significantly higher expression levels observed in obese males (**Table 5**). Additionally, *NPY4R* displayed a significantly opposite pattern in the sex comparisons - upregulated expression in the IOF comparison and downregulated expression in the IOM comparison (**Fig. 2A**).

### Transcriptomic profiles of sex differences in type 2 diabetes

The study of sex differences in the impact of T2D (SDID) did not reveal significant differences in gene expression. We compared the expression profiles of the IOM and IOF comparisons to extract meaningful information from individual sex comparisons (**Fig. 3**). We analyzed 20,161 genes per comparison, with 66 genes exhibiting significantly decreased gene expression in both females and males, and 49 genes exhibiting significantly increased expression in the same comparisons. We did not find any significantly affected genes with an opposite pattern between sexes (i.e., increased in IOM and decreased in IOF or vice versa); however, specific genes displayed discordant patterns among the significant genes in one group, even though the results did not reach significance in the other group (**Fig. 3**).

We identified 63 genes with significantly decreased and 56 genes with significantly increased expression levels in the IOF comparison. These genes displayed an opposite pattern in the IOM comparison, although these results did not reach significance. 5/63 genes had previous associations with obesity, seven with T2D, and five with both disorders, according to information extracted from OpenTargets (**Table 5**) (including *GFAP* and *RENT* among these genes). Of the 56 genes, eight had previous associations with obesity, two with T2D, and two with both disorders, with the *VAX2* gene representing a notable example.

We also observed nine significantly downregulated and eight significantly upregulated genes in the IOM comparison; however, we did not observe a significant pattern in the IOF comparison. Among the significantly affected genes in the IOM comparison, only *ZDHHC19* has a previous association with obesity, according to information extracted from OpenTargets (**Table 5**).

### Functional patterns in obesity and type 2 diabetes

We conducted a GSEA on the gene patterns in each comparison using BP GO terms to provide a functional and integrative perspective on obesity and T2D. Using an FDR < 0.05 cut-off, we identified 2,962 (IO), 2,719 (IOM), 1,808 (IOF), and 31 (SDIO) significantly altered processes in obesity, and 756 (ID), 792 (IDM), 536 (IDF), and 48 (SDID) significant altered processes in T2D.

In the context of obesity, significant functions with the greatest magnitude of change in IOF indicated enrichment in processes such as “*Immunoglobulin-mediated immune response*,” “*B cell-mediated immunity*,” and “*Phagocytosis, engulfment*,” which are associated with antigen recognition and immune response. These functions also become significantly enriched in IOM, albeit with a lower logOR, leading to significant differences in SDIO. In comparison, the most enriched functions in IOM included “*Monocyte chemotaxis*,” “*Antigen processing and presentation of exogenous antigen*,” and “*Antigen processing and presentation of peptide antigen via MHC class II*,” biological processes which are equally enriched in IOF (**Fig. 4A**).

**Fig. 4.**
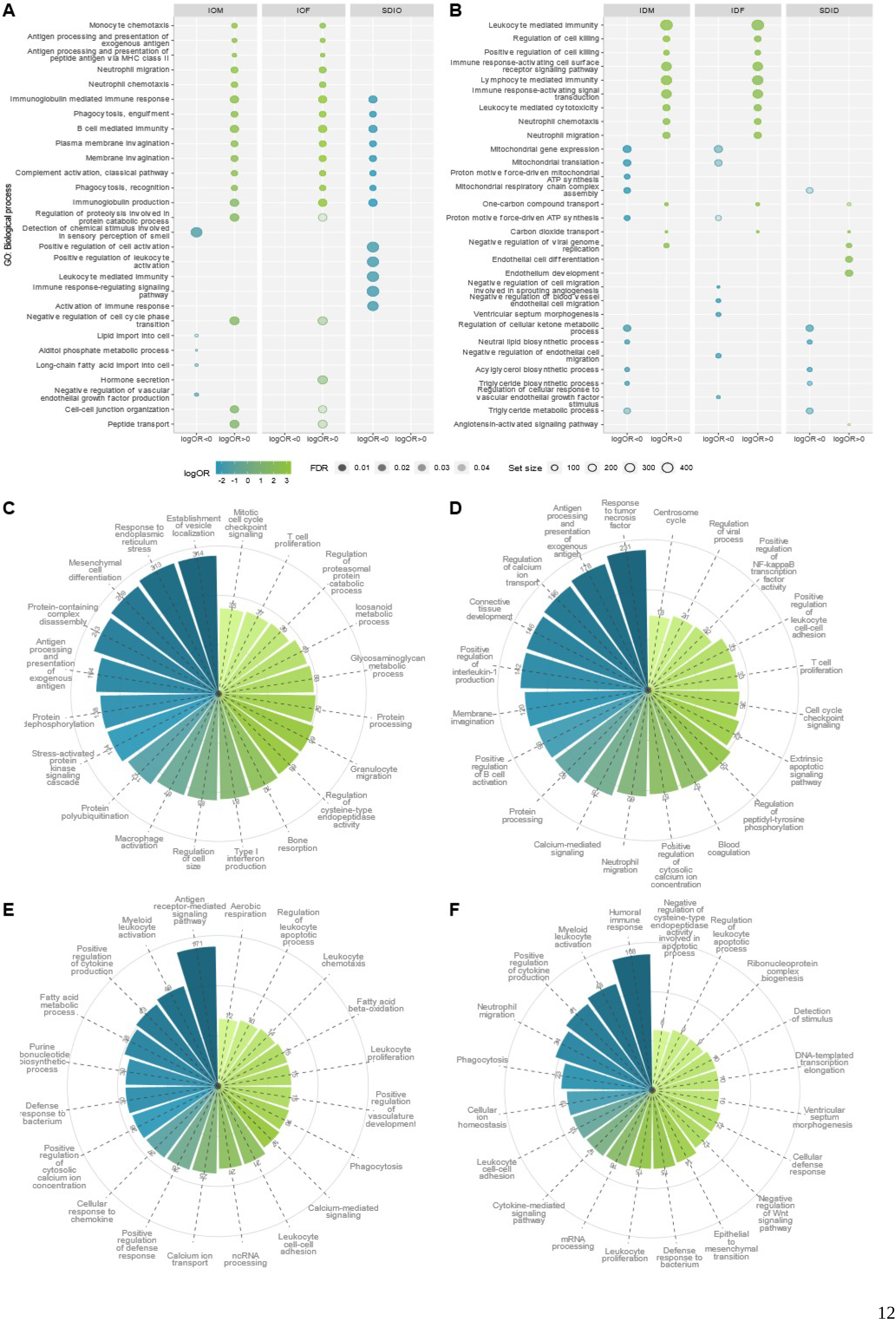
Gene Set Enrichment Analysis Results for Biological Processes Gene Ontology. (**A** and **B**) The processes identified in GSEA for (**A**) obesity and (**B**) T2D have been ordered based on logOR; dot plots represent the top five over- and under-expressed terms per comparison. (**C-F**) The color indicates the effect’s magnitude and direction, while transparency corresponds to the FDR (lower significance results in less opacity). The circle bar size represents the number of genes associated with each enriched term. Graphical summary of the critical functions obtained in the (**C**) IOM, (**D**) IOF, (**E**) IDM, and (**F**) IDF comparisons. BP GO terms were grouped based on distance using the REVIGO methodology. The top twenty groups with the highest number of associated significant BP GO terms are represented, along with the number of processes associated and the name assigned to each after analysis. IOM: impact of obesity in males; IOF: impact of obesity in females; IDM: impact of T2D in males; IDF: impact of T2D in females.

The thirty-one significantly enriched functions in the SDIO comparison were all related to the activation and regulation of the immune system and inflammation, displaying a negative logOR denoting the enrichment of these functions in females in the development of obesity compared to males (**Fig. 4A**).

Notably, the IOM comparison also revealed enriched biological processes with a negative logOR associated with lipid transport and metabolism, including “*Lipid import into the cell*,” “*Alditol phosphate metabolic process*,” and “*Long-chain fatty acid import into the cell*.”

We grouped enriched terms based on distance due to their considerable number and hierarchical structure. In parallel with the role of immunity, we observed an increase in stress-related processes and the role of calcium in the development of obesity in both sexes. These functions are grouped in IOM under the terms “*Establishment of vesicle localization*,” “*Response to endoplasmic reticulum stress*,” “*Stress-activated protein kinase signaling cascade*,” and “*Regulation of cell size*,” totaling 627 terms (**Fig. 4C**). Similarly, in the IOF comparison, the terms “*Calcium-mediated signaling*,” “*Positive regulation of cytosolic calcium ion concentration*,” and “*Regulation of calcium ion transport*” displayed enrichment, with a total of 291 functions (**Fig. 4D**). In addition, in males, the term “P*rotein-containing complex disassembly*” highlights processes related to mitochondrial damage, including those associated with mitochondria and the electron transport chain; in comparison, the number of functions associated with mitochondrial processes in females is negligible.

In the context of T2D, results from both sexes highlight the relevant role of immunity (**Fig. 4B**), consistent with the obesity results; however, the deregulation of biological processes such as “*Regulation of cellular ketone metabolic process*,” “*Neutral lipid biosynthetic process*,” “*Acylglycerol biosynthetic process*,” “*Triglyceride biosynthetic process*,” “*Triglyceride metabolic process*” and “*Mitochondrial respiratory chain complex assembly*” led to substantial changes in the SDID comparison.

Examination of the functional profiles obtained in IDM (**Fig. 4E**) and IDF (**Fig. 4F**) again highlights the role of fatty acid metabolism in obese males, which encompasses over fifty biological processes under the terms “*Fatty acid metabolic process*” or “*Fatty acid beta-oxidation*.” Conversely, T2D females exhibit evidence of fourteen enriched biological processes related to the negative regulation of the Wnt signaling pathway. Based on these results, we compiled 164 genes associated with this pathway - of which fifteen displayed a significant decrease in IDF, while only *PRKN* displayed significant changes in IDM (**Table 5**).

### SDIO and T2D *in silico* signatures experimental validation in obese subjects

To validate *in silico* results, we proceeded to analyze RNA from SAT obese and T2D obese patients by quantitative PCR (qPCR) (Fig. 5). We first selected potential gene expression biomarkers from the SDIO *in-silico* results (Table 4) to validate differential gene expression between obese males and females. Overall, the expression of the *SPATA18, KREMEN1, NPY4R*, and *PRM3* genes confirmed the *in silico* results meta-analysis (**Fig. 5A**).

**Table 4.** Significantly affected genes identified in the obesity meta-analysis. The significant genes are grouped by comparison and logFC sign. The genes have been ordered based on the FDR and the absolute value of the effect size, in presence of a large set of significant genes only the first 20 are shown, as well as some relevant and significant genes for further interpretation. The genes are highlighted according to their association with obesity (bold), T2D (underlined), or both, based on the information extracted from Open Targets Platform.

| Comparison | logFC > 0 | logFC < 0 |
| --- | --- | --- |
| IO | WDR41, **FCGR2B**, LAMB2, SERPINH1, RNASE6, AGTRAP, AP2S1, HCLS1, IRF8, <u>TYK2</u>, ALOX5AP, TMEM109, CAPZB, IL10RA, **VSIG4**, **PLA2G7**, HYCC1, GPR137B, **ACP5**, <u>NOP2</u>, (+410) | TEX51, MTHFS, <u>INSR</u>, **TMPRSS6**, KYAT3, **MAOA**, <u>SLC6A5</u>, ECPAS, URAHP, LOC105373554, MPDZ, RPLP2, ARX, PPP4R1L, LINC02833, ACADSB, CACHD1, DCST1-AS1, ST6GALNAC6, APTR, (+330, including: LINC01208) |
| IOM | ANXA11, **CCND1**, ACTN1, DDX41, YKT6, LOC101929398, <u>STAT1</u>, CPVL, IGLC2, **PLA2G7**, **ANXA5**, S100A11, TAPBPL, ITGB5, PLXNB2, SEC31A, CAPZB, GPR137B, **ACP5**, LAMB2, (+257, including: SLAMF8, **SPATA18**) | <u>CRH</u>, LINC01208, MCCC1, TMOD1, HADHB, **ECHS1**, **HK2**, USP32P2, **MAMLD1**, TEX51, CASD1, COPG2IT1, **MAOA**, <u>PKP2</u>, SALL2, LINC01788, PCDHA5, RPS3AP20, TAC4, TRMT9B, (+117, including: <u>INSR</u>, **NPY4R**) |
| IOF | HCLS1, **CD163**, **CHTOP**, <u>LYN</u>, ARHGAP17, PRSS23, **PPRC1**, RNASE6, **PLA2G7**, IGHG2, PLA2G4C, **ABCC5**, IGHV3-23, GAB3, PPP4R1, <u>PUS1</u>, BCYRN1, <u>SEPTIN9</u>, FKBP15, **FCGR2B**, (+127, including: **NPY4R**) | IGHD6-6, MTHFS, TRGV5, STAU2-AS1, **RAD21**, ISCU, RSPH10B2, **NDUFA2**, OPHN1, ST6GALNAC6, POLR3GL, GKAP1, TEX51, LINC02266, DCAF12L1, **RBL2**, **EIF4EBP2**, PRDM12, RTP1, NKAIN3, (+143, including: <u>INSR</u>, LOC101928371, MIF-AS1, COPB2-DT, LINC02126, MID1IP1) |
| SDIO | LOC101928371, MIF-AS1, SLAMF8, AMER1, COPB2-DT, **SPATA18**, LINC02126, ANXA11, MID1IP1, ELK1 | **NPY4R**, **KREMEN1**, LINC01208, ZSCAN5A, UBASH3B, **ITIH4**, TPSG1, <u>PRM3</u>, **MAEA** |

**Table 5.**
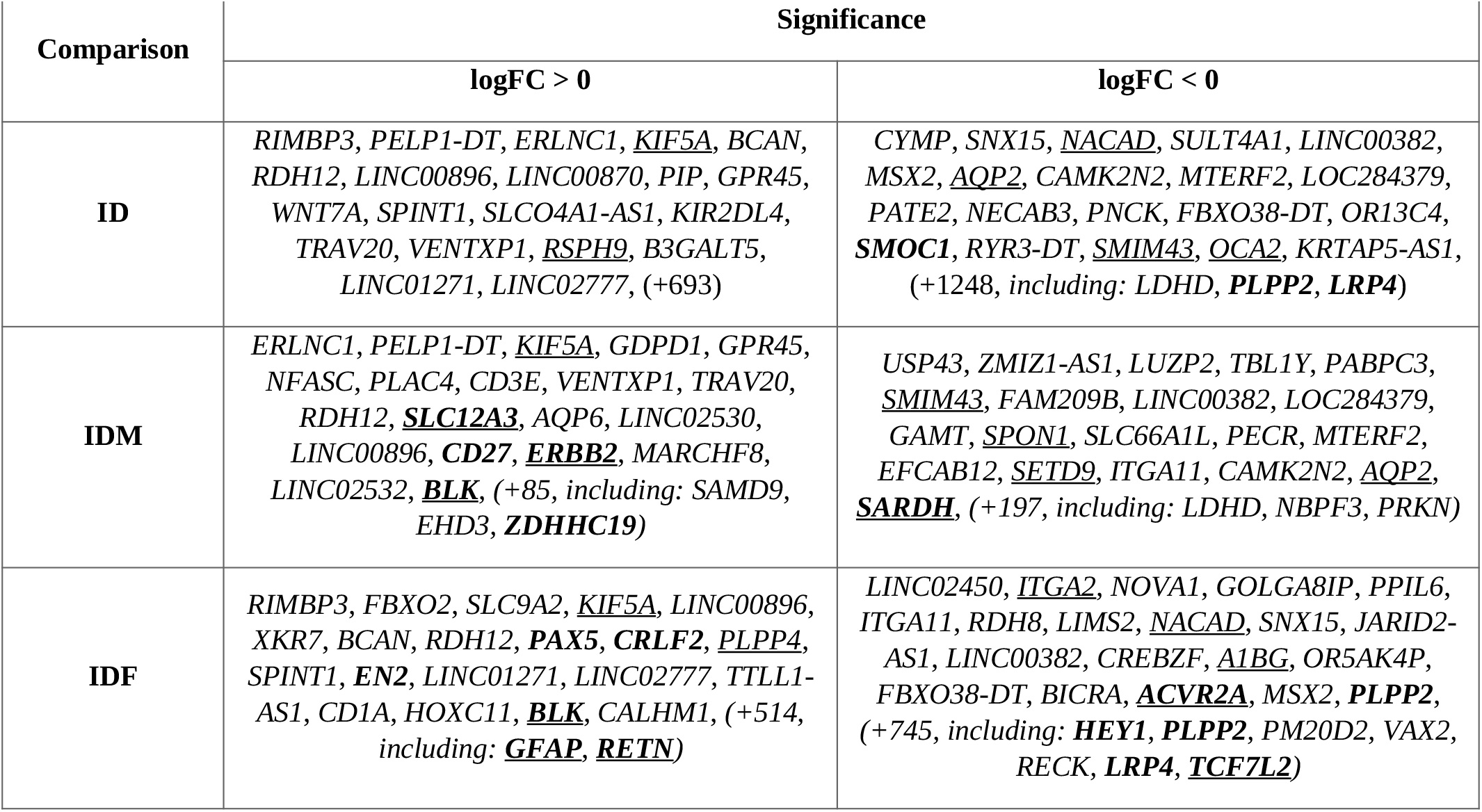
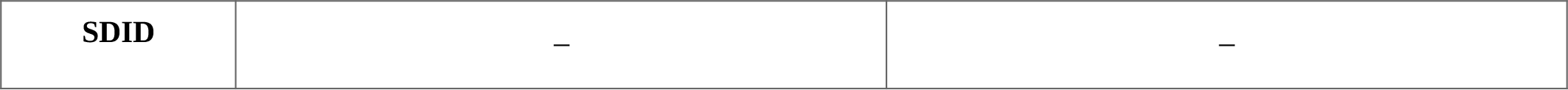
Significantly affected genes identified in the type 2 diabetes meta-analysis. The significant genes are grouped by comparison and logFC sign. The genes have been ordered based on the FDR and the absolute value of the effect size, in the presence of a large set of significant genes only the first 20 are shown, as well as some relevant and significant genes for further interpretation. The genes are highlighted according to their association with obesity (bold), T2D (underlined), or both, based on the information extracted from Open Targets Platform.

**Fig. 5.**
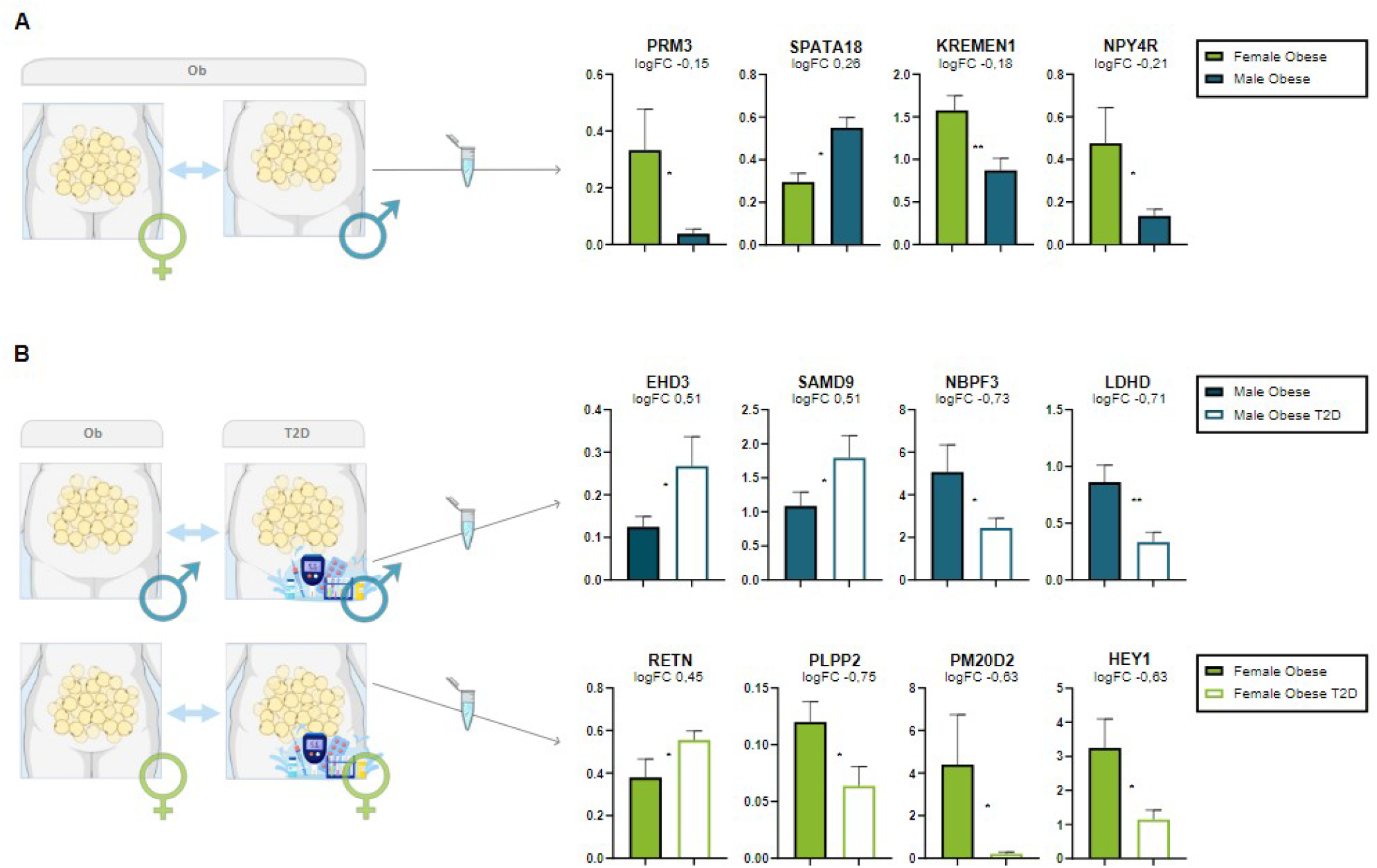
Gene expression-mediated validation of the *in-silico* results. Relative gene expression analysis in human SAT samples from obese and obese T2D males and females by qPCR. After obtaining informed consent, SAT samples were obtained during surgery, and RNA was extracted and used for real-time PCR. (**A**) Relative gene expression analysis for obesity in male and female obese samples. (**B**) Relative gene expression analysis for T2D in males (upper panel) and females (lower panel). In *silico-analyses, fold-change* results are shown for each gene. Each experimental group is represented by an n=8. Student’s t-test applied for significance: (*) p-value<0.05, (**) p-value<0.01. Created with BioRender.com.

We then employed qPCR analysis to confirm a T2D signature specific for males and females as revealed by differential expression profile between T2D and obese males (*SAMD9, NBPF3, LDHD*, and *EHD3*) and females (*RETN, HEY1, PLPP2*, and *PM20D2*) (**Fig. 5B**). We selected genes according to *in silico* significant differences (Table 5), playing key metabolic roles, and found to be referenced for its expression in SAT in the Genotype-Tissue Expression (GTEx) Portal.

In all cases, to validate SDIO, and male and female T2D signatures with subcutaneous adipose tissue samples, we selected *RPL19* as the reference gene due to its previously demonstrated stability and consistency in WAT when considering sex as a biological variable [86, 87].

## Discussion

Although we know of significant differences between sexes in the prevalence, development, and pathophysiology of obesity and T2D, their molecular basis remains unclear. Understanding the impact of sex-specific differences in disease remains critical to improving outcomes. To the best of our knowledge, we conducted the first systematic review and meta-analysis on WAT with sex as a central perspective, revealing gene biomarkers and functions involved in obesity and T2D progression using publicly available data.

We used an *in-silico* approach, analyzing transcriptomic studies that included WAT samples from patients of both sexes with obesity and T2D and control patients. We analyzed data via eight meta-analyses and subsequent functional analyses, identifying common patterns and biomarkers specific to each disorder and the more significant or exclusive involvement of specific biological processes in the development of obesity and T2D in SAT in relation to sex.

### Transcriptomic landscape of sex differences in obesity

Our meta-analysis identified 1,185 potential unique biomarkers involved in weight gain and obesity across comparisons. Of these, 18.8% had previous associations with obesity,18.5% with T2D, and 8.4% with both, according to data extracted from OpenTargets (**Table 4**) adding robustness and novelty to our findings. Further analysis revealed twenty-six common genes between the sexes (**Fig. 2A**), including *INSR*, which plays a significant role in insulin resistance and T2D development/progression [12].

In terms of sex differences in the impact of obesity, we identified nineteen potential biomarker genes (**Fig. 2**); of note, five represent long non-coding RNAs (*LOC101928371, LINC02126, LINC01208, COPB2-DT*, and *MIF-AS1*) [88–92]. Among these, *MIF-AS1* has a pro-inflammatory role in obesity and is involved in the development of insulin resistance [106]. Six additional genes (*SLAMF8, ZSCAN5A, UBASH3B, TPSG1, MAEA*, and *ANXA11*) participate in the activation and differentiation of macrophages, lymphocytes, and mast cells [93] and the production of pro-inflammatory cytokines [94, 95], and its implications in T2D susceptibility [96], progression [97], and complications [98] have been widely reported. Our results also highlighted other diabetes-associated genes - *ITIH4* and *PRM3* [99]- which were shown to be overexpressed females compared to males during the experimental validation in SAT samples (**Fig. 5A**).

*SPATA18* represents a critical gene involved in the induction of mitophagy and the regulation of mitochondrial quality in response to DNA damage [100]. Our study revealed sex-specific *SPATA18* expression, with upregulated expression in obese males (**Fig. 2**; **Fig. 5A**) indicating higher susceptibility to mitochondrial damage and the development of T2D. Moreover, *ELK1* and *MID1IP1* exhibited a similar pattern in obese males, supporting a higher risk of T2D development in obese males. *ELK1* encodes a transcription factor essential for T2D development that becomes activated by phosphorylation via the MAPK/ERK1/2 pathway, promoting adipogenesis and insulin resistance in WAT [101, 102]; meanwhile, increased *MID1IP1* expression has been associated with lipogenesis and fatty acid accumulation [103].

The Wnt β-catenin pathway regulates metabolism and homeostasis, inhibiting adipogenesis and increasing insulin sensitivity via negative regulation of *PPAR-γ* and *GSK3β* [12]. As genes encoding antagonists of this pathway [104, 105], *KREMEN1* exhibits increased expression in obese females and decreased expression in obese males (**Fig. 2**; **Fig. 5A**), while *AMER1* displays the opposite trend. The altered transcriptional regulation of these two genes could imply the inhibition of the Wnt β-catenin pathway as one of the primary regulators in the development of T2D in obese patients, regulated by complementary sex-dependent mechanisms.

*NPY4R*, has emerged as a sex-specific biomarker that may contribute to diet-induced obesity in females [106, 107], although this gene plays an unknown role in males. Our study revealed the significant upregulation of *NPY4R* expression in obese females and downregulation in obese males (**Fig. 2**; **Fig. 5A**). Previous studies associated increased *NPY4R* expression with an orexigenic effect and a positive correlation with body mass index and abdominal circumference only in females [106, 107]. However, this gene has never been studied from a sex-specific perspective before.

Functionally, we observed the enrichment of inflammatory processes in obese patients of both sexes, but more significantly in females (**Fig. 4**); furthermore, we observed a sex-neutral enrichment in processes related to stress and the role of calcium in the development of obesity. Meanwhile, obese males exhibited processes related to mitochondrial damage, which supports our gene-level findings.

### Transcriptomic landscape of sex differences in type 2 diabetes

The meta-analysis conducted on T2D samples led to the identification of 2,542 potential biomarkers. 15.2% possessed previous associations with obesity, 16.2% with T2D, and 5.6% with both conditions (**Table 5**). Individual comparisons by sex, IDM, and IDF revealed sex-specific potential biomarkers absent in the global comparison, with a higher number in females due to their greater representation in the included studies. Although the methodology and the limited number of studies and samples prevented the identification of differentially expressed genes in the study of sex differences in diabetes (SDID), comparing profiles by sex allowed us to draw some interesting conclusions.

*FAM83E* and *RASAL1* represent potential biomarkers overexpressed in T2D males; both genes participate in the Ras/MAPK signaling pathway and associate with body weight, T2D, and complications such as diabetic nephropathies [108–110]. *EHD3*, which participates in protein biosynthesis and transport [111], and *SAMD9*, which encodes a plasma protein involved in inflammatory response through TNF-alpha [112], have associations with diabetes pathology and complications. These genes are generally expressed in several adipose tissue cell types, and their overexpression in male T2D has been assessed for the first time in male human SAT (**Fig. 3**; **Fig. 5B**). Significantly, we also detected the overexpression of *NBPF3*, expressed in spermatids in testis, in male obese samples compared to male T2D samples, suggesting a potential connection between T2D and fertility (**Fig. 3**; **Fig. 5B**).

The *GFAP* and *RETN* genes displayed significant overexpression, and the *VAX2* gene showed significant downregulation in female T2D samples, indicating their potential as female-specific biomarker genes. Previous studies investigated the role of *GFAP* in the development of T2D, revealing notably higher gene expression in females [113]. *RETN* gene polymorphisms have been associated with an increased risk of T2D [110, 114], with significant differences observed between sexes [115]. Here, we validated *RETN* overexpression in female T2D SAT samples (**Fig. 5B**); and *PLPP2, PM20D2*, and *HEY1* overexpression in female obese samples compared to T2D counterparts. These markers participate in adipose tissue metabolism in lipids (*PLPP2)*, proteins (*PM20D2)*, and at the transcriptional level (*HEY1)* (**Fig. 5B**); furthermore, their expression becomes enriched in several cell types from SAT and VAT according to GeneCards and Human Protein Atlas [116, 117].

At the functional level, both sexes maintain a significant role of immunity in relation to obesity; however, the SDID comparison highlighted a dysregulation of biological processes related to fatty acid biosynthesis and β-oxidation, which is mainly significant in T2D males. In addition, we observed greater dysregulation of mitochondrial processes in T2D samples, as found with *LDHD* (**Fig. 3**; **Fig. 5B**), leading to significant differences between sexes (**Fig. 4B**): of note, these findings agree with the results of *SPATA18* observed in obese patient samples.

Examining functional profiles by sex (**Fig. 4E-F**) revealed the presence of fourteen significant functions associated with the Wnt signaling pathway in T2D females, which do not display significance in males. These biological processes are linked to fifteen genes, including key regulators of both canonical and non-canonical Wnt pathways. *RECK* encodes a protein that acts as a Wnt7-specific coactivator of the canonical Wnt signaling pathway by decoding Wnt ligands. *LRP4* encodes the low-density lipoprotein receptor-related protein 4, which regulates the WNT/ß-catenin pathway and whose expression levels have been previously associated with metabolism, insulin response, and diabetes [24–27]. *VAX2* encodes a dominant-negative Wnt antagonist that regulates the expression of Wnt signaling antagonists, including the expression of a truncated *TCF7L2* isoform and *TCF7L2*, a critical transcriptional effector of the Wnt/β-catenin signaling pathway. Other involved genes include *TRABD2B, TMEM64, APCDD1L, TCF7L1, ARHGEF19, WLS, BCL9, TBX18, DRD2, GSK3A* and *NFATC1*.

Additionally, the sex-specific biomarker for obesity, *KREMEN1*, exhibits inconsistent regulation in T2D samples from both sexes, with non-significantly increased expression in females and decreased expression in males. These findings emphasize the significant role of the Wnt signaling pathway in the development of obesity-related T2D, particularly in females, and its strong association with sex, as observed in obesity.

### Strengths and limitations

Sex-related differences have been documented in the clinical and pathophysiological aspects of metabolic diseases and disorders, including obesity and T2D [37, 38, 47]; however, sex is often perceived more as a bias rather than a biological variable in biomedical research. Encouragingly, the National Institutes of Health [118] and the European Union [119, 120] have recently recognized sex as a necessary biological variable in research. Incorporating this perspective is critical for identifying sex differences and generating more accurate, clinically relevant results.

Our study has been able to address sex as a biological variable in the development of T2D based on obesity in human SAT, representing a significant advance in understanding male and female differences.

We focused our systematic review on subcutaneous and vascular white adipose tissues of organisms Mus musculus and Homo sapiens. However, the lack of sex-specific data on human VAT and mouse WAT was our first limitation, leading to the exclusion of a significant proportion of samples studies and restricting the analysis to human SAT.

Despite efforts to publish open data under the FAIR (Findable, Accessible, Interoperable, Reusable) principles [121], data availability remains limited. During our search, 24% of identified publications were not retrievable due to restricted access or missing datasets. This, along with the lack of homogeneity in the clinical and phenotypic data, prevented the integration of variables like age, body mass index, or insulin resistance index, which could have impacted disease variability.

These limitations restricted the number of studies and samples analyzed, with a higher representation of females. This imbalance led to a loss of statistical power, especially when studying the impact of T2D.

On the other hand, our methodology enhanced the robustness and reliability of the identified biomarkers by accounting for variability between samples and studies, thereby facilitating their extrapolation to the general population. As a result, both novel and previously known sex-specific biomarkers for obesity and T2D were identified and validated in vitro.

Our meta-analysis revealed common and differential gene expression patterns between sexes in developing obesity and T2D, identifying potential biomarkers. While some of our findings had previously been reported -albeit not typically from a sex perspective-, others represent novel discoveries. These differentially expressed genes offer promising avenues for further research, with implications for disease diagnosis, prognosis, and the design of clinical trials.

### Perspectives and significance

Our meta-analysis identified common and differential gene expression patterns between sexes in developing obesity and T2D, identifying potential biomarkers. While some of our findings had previously been reported (albeit not typically from a sex perspective), others represent novel discoveries. Identifying differentially expressed genes provides new research opportunities to investigate their role in obesity and their potential as sex-specific biomarkers or risk factors for obesity and T2D.

This study provides evidence of male- and female-specific transcriptional signatures in SAT related to the development of obesity and T2D. The prominent role of mitochondria and altered fatty acid metabolism in males may explain their higher predisposition to T2D in obesity. In addition, the involvement of the Wnt pathway, particularly in females, offers new pathways for the exploration and study of potential therapeutic targets.

These findings have implications for disease diagnosis, prognosis, and the design of clinical trials. Furthermore, using publicly available data, our study employs a novel bioinformatics strategy to assess sex-based differences; this highlights the importance of considering sex as a variable when studying metabolic diseases.

## Conclusions

The sex perspective remains largely overlooked in biomedical research. The *in-silico* integration of public data using a meta-analysis methodology enabled the discovery of common patterns and potential sex-specific biomarkers in adipose tissue associated with the development of obesity and T2D, highlighting the importance of sex in metabolic disorders and paving the way for new avenues of research towards personalized medicine.

## Abbreviations

BH: Benjamini & Hochberg
BP: Biological Processes
DGE: Differential Gene Expression Analysis
F: female
FDR: False discovery rate
GEO: Gene Expression Omnibus
GO: Gene Ontology
GSEA: Gene Set Enrichment Analysis
ID: Impact of Diabetes
IDM: Impact of T2D in Males
IDF: Impact of T2D in Females
IO: Impact of Obesity
IOF: Impact of Obesity in Females
IOM: Impact of Obesity in Males
logFC: logarithm 2 of the Fold Change
logOR: logarithm 2 of the Odds Ratio
M: Male
qPCR: quantitative PCR
SAT: Subcutaneous Adipose Tissue
SDIO: Sex-Differential Impact of Obesity
SDID: Sex-Differential Impact of T2D
T2D: Type 2 Diabetes
VAT: Visceral Adipose Tissue
WAT: White Adipose Tissue

## Consent for publication

Not applicable.

## Availability of data and materials

The data and results generated in the various steps of the meta-analyses are freely available on the Metafun-SDT2D platform (http://bioinfo.cipf.es/metafun-SDT2D), and all scripts are available in the GitHub repository https://github.com/Roxyam/T2D-Meta-Analysis.

## Competing interests

The authors declare that they have no competing interests.

## Funding

This research was supported by and partially funded by the Institute of Health Carlos III (project IMPaCT-Data, exp. IMP/00019), co-funded by ERDF, “A way to make Europe”; PID2021-124430OA-I00 funded by MICIU/AEI/10.13039/501100011033 by FEDER, UE; and SAF2017-84708-R grants.

## CRediT authorship contribution statement

**Roxana Andreea Moldovan:** Methodology, Software, Formal analysis, Investigation, Data curation, Writing—original draft preparation, Visualization. **Marta R. Hidalgo**: Methodology, Software, Investigation, Writing—original draft preparation, Writing—review and editing, Visualization, Supervision, Project administration. **Helena Castañé**: Writing—original draft preparation, Writing—review and editing. **Andrea Jiménez-Franco**: Writing—original draft preparation, Writing—review and editing. **Jorge Joven**: Writing—original draft preparation, Writing—review and editing. **Deborah J. Burks**: Writing— original draft preparation, Writing—review and editing, Funding acquisition. **Amparo Galán**: Conceptualization, Methodology, Formal analysis, Investigation, Experiment conduction, Writing—original draft preparation, Writing—review and editing, Supervision, Project administration. **Francisco García-García**: Conceptualization, Methodology, Investigation, Writing—original draft preparation, Writing—review and editing, Supervision, Funding acquisition, Project administration.

## Acknowledgments

The authors thank the Principe Felipe Research Center (CIPF) for providing access to the cluster, co-funded by European Regional Development Funds (FEDER) in the Valencian Community 2014-2020, and Stuart P. Atkinson for critically reviewing/editing the manuscript.

## Notes

http://bioinfo.cipf.es/metafun-SDT2D

https://github.com/Roxyam/T2D-Meta-Analysis

### Competing Interest Statement

The authors have declared no competing interest.

